# Axonal Na^+^ channels detect and transmit levels of input synchrony in local brain circuits

**DOI:** 10.1101/618710

**Authors:** Mickaёl Zbili, Sylvain Rama, Pierre Yger, Yanis Inglebert, Norah Boumedine-Guignon, Laure Fronzaroli-Moliniere, Romain Brette, Michaёl Russier, Dominique Debanne

## Abstract

Sensory processing requires mechanisms of fast coincidence-detection to discriminate synchronous from asynchronous inputs. Spike-threshold adaptation enables such a discrimination but is ineffective in transmitting this information to the network. We show here that presynaptic axonal sodium channels read and transmit precise levels of input synchrony to the postsynaptic cell by modulating the presynaptic action potential (AP) amplitude. As a consequence, synaptic transmission is facilitated at cortical synapses when the presynaptic spike is produced by synchronous inputs. Using dual soma-axon recordings, imaging, and modeling, we show that this facilitation results from enhanced AP amplitude in the axon due to minimized inactivation of axonal sodium-channels. Quantifying local circuit activity and using network modeling, we found that spikes induced by synchronous inputs produced a larger effect on network activity than spikes induced by asynchronous inputs. Therefore, this input-synchrony dependent facilitation (ISF) may constitute a powerful mechanism regulating spike transmission.

---

Higher brain function such as sensory processing or encoding of memory requires biological mechanisms able to discriminate highly synchronized inputs from weakly synchronized or asynchronous inputs with a temporal accuracy in the millisecond range^1–4^. This property is classically thought to be achieved by mechanisms of coincidence detection occurring at the postsynaptic side such as activation of NMDA receptors at glutamatergic synapses^5^, dendritic amplification of synaptic responses^6^, and spike-threshold dynamics^7^. Spike-threshold adaptation represents the most accurate mechanism known for detection of fast input synchrony in central neurons^7–10^. The spike-threshold recorded in cortical neurons *in vivo* depends on the rate of depolarization and the nature of the stimulus^7, 11^. For example, in the barrel cortex, the spike-threshold is hyperpolarized for the preferred stimulus and depolarized for the non-preferred stimulus^11^. Theoretical work has underlined the importance of Na^+^ channel inactivation in the adaptation of spike-threshold^9, 12^. But it is not known whether the readout of input synchrony by adaptation of spike-threshold can be transmitted within the network. This amounts to asking the question: does an action potential produced by synchronous inputs convey more information than a spike produced by asynchronous inputs? Axonal ion channels have been shown to determine presynaptic spike waveform and hence, synaptic transmission in a context-dependent manner that corresponds to analog-digital modulation of synaptic strength^13–18^. We therefore examined whether levels of input synchrony could be read and transmitted through the network by axonal channels.

We show here that axonal sodium channels detect levels of input synchrony with millisecond precision and transmit this information to the network through modulation of action potential amplitude and presynaptic release. In fact, synaptic transmission is facilitated at cortical connections when the presynaptic action potential is produced by synchronous inputs. Using dual soma-axon recordings, calcium imaging and computer modeling, we show that this facilitation results from the enhancement of axonal spike amplitude caused by minimized inactivation of axonal sodium channels. In addition, we reveal that synchronous input-mediated spikes provoke a larger increase in local circuit activity than asynchronous input-mediated spikes. This suggests that this form of context-dependent enhancement of synaptic transmission may represent a robust way for information coding.

## Results

### Input-synchrony-dependent facilitation

Pairs of monosynaptically connected L5 pyramidal neurons were recorded in acute slices of P13-20 rats. The effect of synchronous activation on synaptic transmission was tested by injecting AMPA-like excitatory pre-synaptic potentials (EPreSPs) into the presynaptic neuron using dynamic-clamp. Postsynaptic responses were found to be larger when the presynaptic action potential (AP) was produced by highly synchronized inputs (0 ms delay between inputs) than when elicited by intermediate (5 ms delay) or weakly synchronous (10 ms delay) inputs (**Fig. 1a**; 140 ± 14%, n = 8; Wilcoxon-test, p < 0.01). Therefore, the synchrony level of the inputs determines output synaptic strength. We named this phenomenon input synchrony-dependent facilitation (ISF). As previously reported, the AP threshold of the presynaptic neuron was found to be lower as EPreSP synchrony increased (**Supplementary Fig. 1**).

**Fig. 1.**
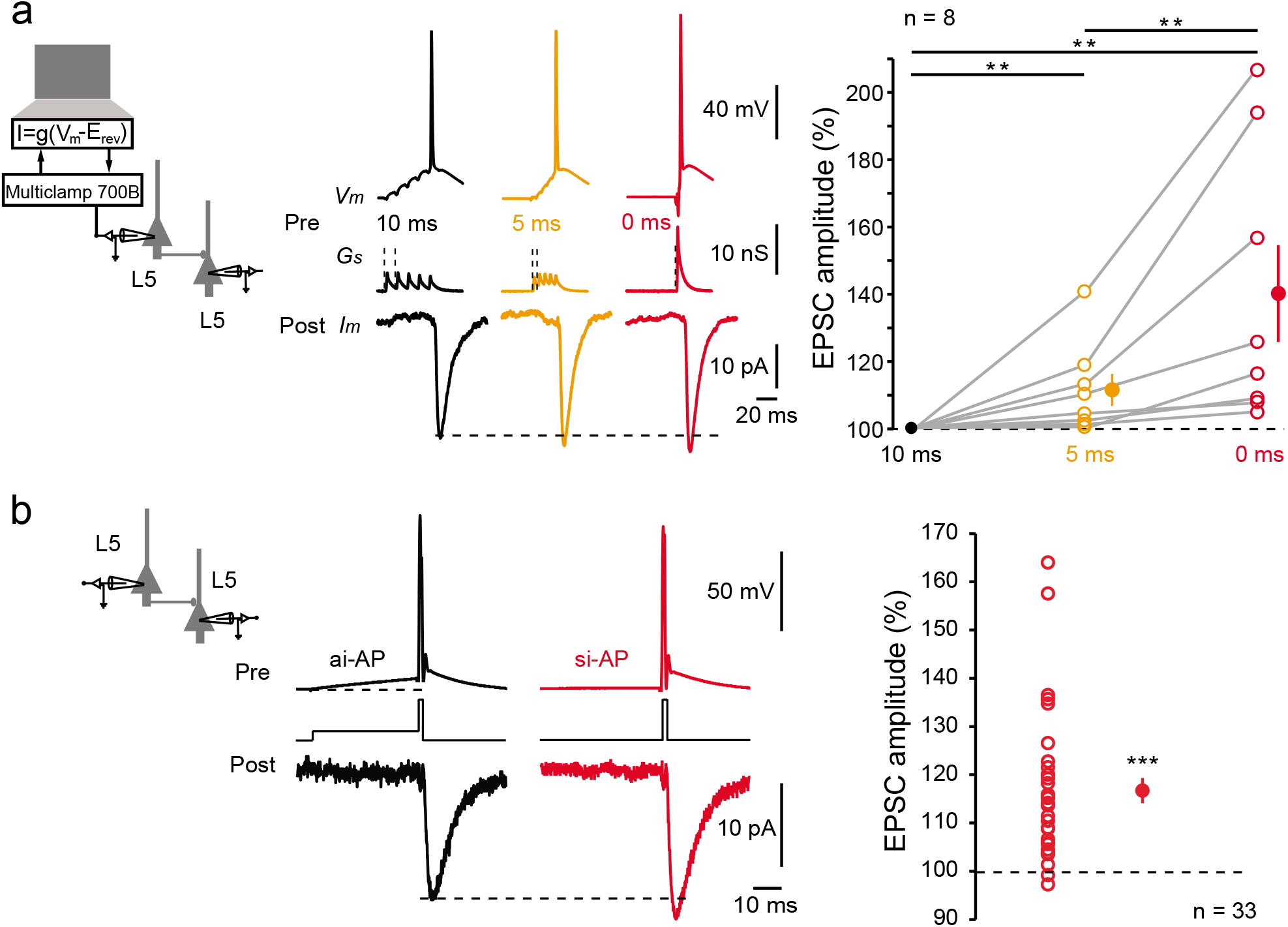
Input-synchrony-dependent facilitation (ISF) at L5-L5 synapses. **a**, Induction of ISF with physiological stimuli. Left, schematic representation of the system used to inject a dynamic current mimicking 5 glutamatergic inputs into the presynaptic neuron. Middle, examples of electrophysiological recordings from a connected pair of L5 neurons. The 5 EPreSGs were injected with a delay of 10 ms (black traces), 5 ms (orange traces) or synchronously (red traces). Note the increase in the postsynaptic response amplitude with the increase in EPreSGs synchrony. Right, statistics (open circles, individual pairs, closed circles, averages). **b**, Induction of ISF with current pulses. The presynaptic AP was induced by a brief step of current with or without preceding 50 ms pre-pulse to mimic weakly synchronized or highly synchronized inputs, respectively. Note the increased EPSC amplitude induced by a presynaptic AP induced by a synchronous-like input (si-AP) compared to that produced by an asynchronous-like input (ai-AP). Right, plot of input synchrony-dependent facilitation observed in 33 cell pairs.

ISF could also be induced by a simple protocol based on two ways of inducing the presynaptic AP (i.e. directly from resting potential to mimic synchronous inputs or after a 50 ms depolarizing pre-pulse to mimic asynchronous inputs). Consistent with previous findings, APs emitted by a synchronous-like input (si-APs) produced larger postsynaptic responses than APs emitted by asynchronous-like input (ai-APs) (117 ± 3%, n = 33; Wilcoxon-test, p < 0.001; **Fig. 1b**). During ISF, the paired-pulse ratio (PPR) diminished (from 64 ± 4% to 47 ± 3%, n = 24; Wilcoxon-test, p < 0.001; **Fig. 2a**), and the CV^−2^ of EPSC amplitude increased (181 ± 22% of the control, n = 26; Wilcoxon test, p < 0.001; **Fig. 2b**), indicating the presynaptic origin of ISF. Magnitude of ISF was not related to age (**Supplementary Fig. 2**) but followed an inverse function of synaptic strength (**Supplementary Fig. 2**). ISF was also observed at local excitatory synapses in area CA3 of the hippocampus (117 ± 3% of the control EPSC, n = 20; **Supplementary Fig. 3**), indicating that ISF may be a general feature in excitatory brain circuits.

**Fig. 2.**
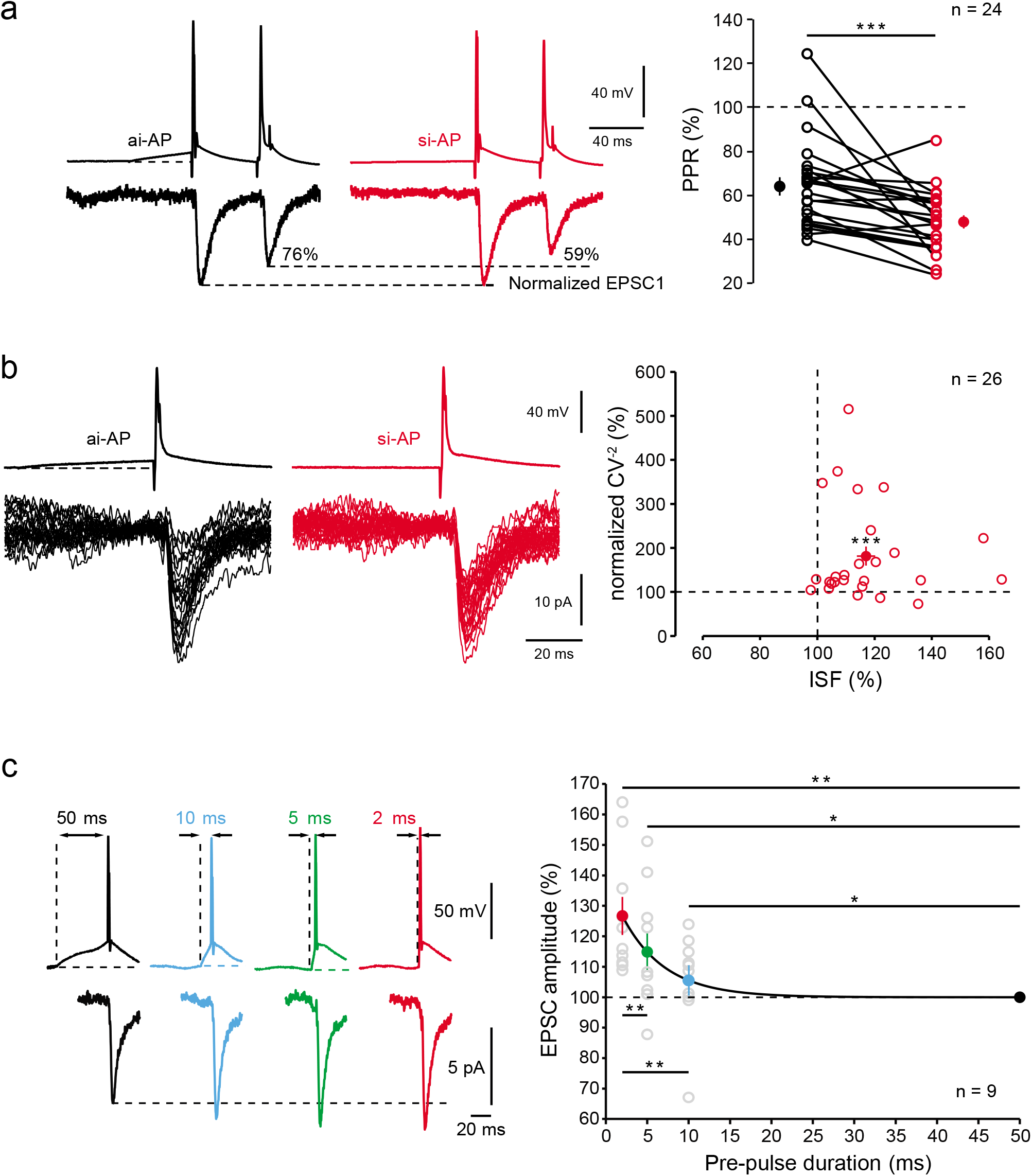
Properties of ISF. **a**, Reduced paired-pulse ratio (PPR) after ISF. Left traces, PPR with an asynchronous-like weak (black) and synchronous-like (red) input. Note the decrease in PPR (here from 76% to 59%). Right, group data. **b**, Reduced EPSC variability during ISF. Left, examples of 30 EPSCs evoked by an asynchronous-like (back) or synchronous-like (red) input. Right, plot of normalized CV^−2^ as a function of ISF. **c**, Time-course of ISF. Presynaptic spikes were evoked with pre-pulse duration (PD) ranging from 50 to 2 ms. Note that evoked postsynaptic responses increased when the PD decreased. Right, plot of EPSC amplitude normalized to that produced by a PD of 50 ms (black line, exponential regression *y* = 39.6 ∗ e^−x/τ^ + 99.98 with *τ* = 5.1 ms; R^2^ = 0.3).

We next determined the time-course of ISF by measuring ISF magnitude as a function of the pre-pulse duration (PD). Presynaptic spikes were induced following depolarizations ranging between 50 and 2 ms (corresponding to input synchrony increase) and the resulting postsynaptic responses were recorded. ISF magnitude followed an exponential decay with increasing PD (time-constant of 5.1 ms; **Fig. 2c**). Importantly, synaptic facilitation observed with a PD of 2 ms was found to be significantly larger than that observed for a PD of 5 ms (Wilcoxon-test, p < 0.01). We conclude that ISF is highly time discriminative.

### Mechanisms of ISF: modulation of axonal spike amplitude

Since ISF has a presynaptic origin, we examined whether ISF was associated with an elevation in presynaptic Ca^2+^ influx. L5 neurons were recorded with a pipette filled with 50 μM Alexa 594 and 250 μM Fluo-4, and spike-evoked Ca^2+^ signals were measured in putative *en passant* boutons (**Fig. 3a**). The amplitude of the spike-evoked Ca^2+^ transient was increased when the spike was evoked by a synchronous-like input (144 ± 8%, n = 6, Wilcoxon-test, p < 0.05; **Fig. 3a**). Interestingly, the ratio between si-AP-evoked Ca^2+^ entry and ai-AP evoked Ca^2+^ entry declined with the distance from the soma with a space constant of 232 μm (**Fig. 3b**), probably due to the decremental propagation of the subthreshold depolarization along the axon (with a space constant of 360 μm; **Supplementary Fig. 4**).

**Fig. 3.**
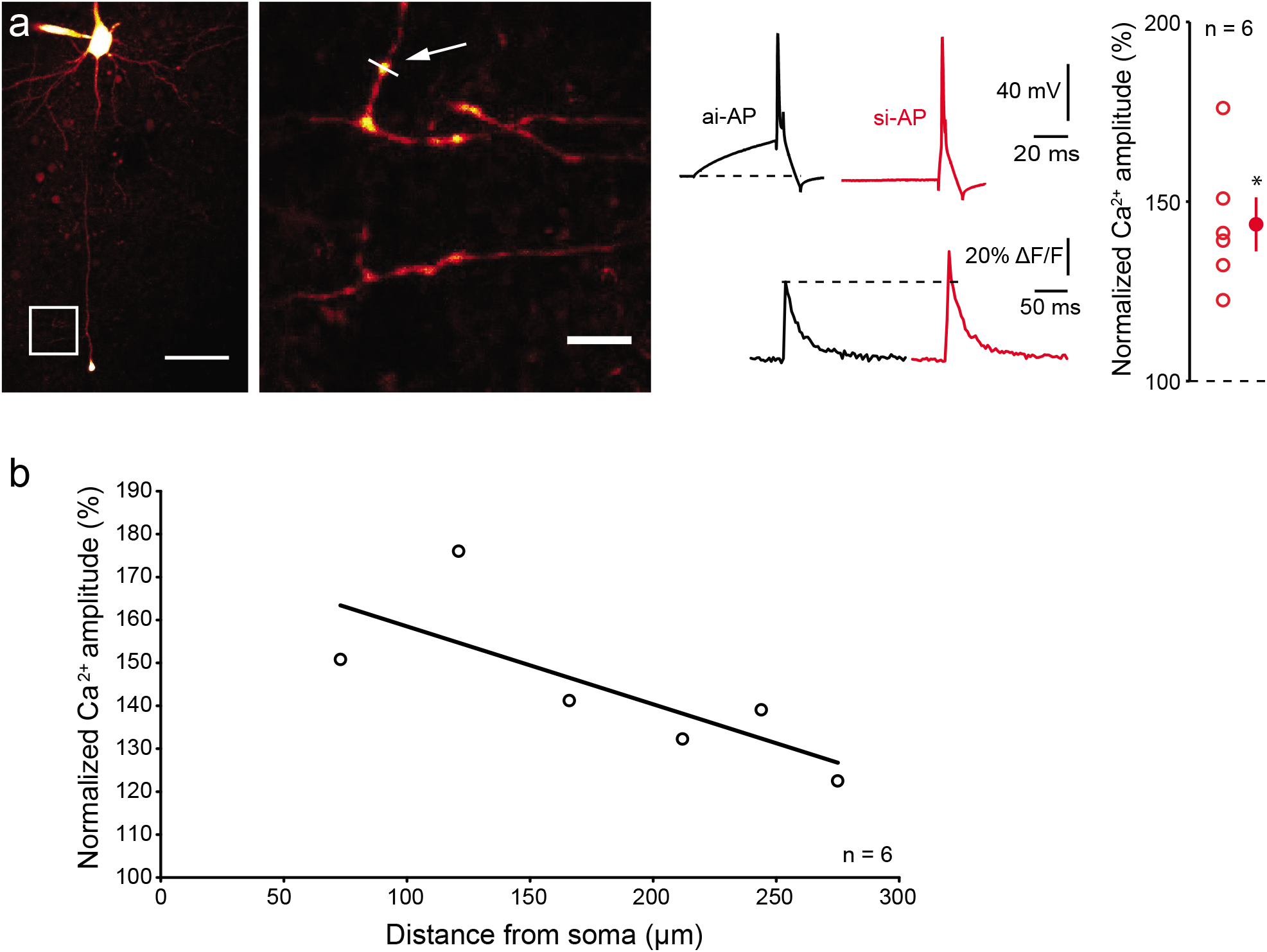
Ca^2+^ imaging. **a**, Left, ensemble view and detail of axonal arborization of an L5 pyramidal neuron filled with Alexa 594 and Fluo-4. The white arrow indicates the line scan made at ~200 μm from the cell body. Scale bars, 50 μm and 2 μm. Middle traces, simultaneous electrophysiological (up) and Ca^2+^ imaging (bottom) recordings during ISF. Note the increase in Ca^2+^ fluorescence when the spike is evoked by a synchronous-like input compared with a spike evoked by an asynchronous-like input. Right, normalized amplitude of the spike-evoked Ca^2+^ signal. **b**, Attenuation of the enhanced spike-evoked Ca^2+^ influx produced by si-APs along the axon. Plot of the normalized amplitude of Ca^2+^ signal as a function of the distance from the soma. The data have been fitted with an exponential regression (*y* = 89.6 ∗ *e^−x/λ^* + 100, with λ = 232 μm, R^2^ = 0.65).

To examine the presynaptic spike shape, L5 neurons were recorded simultaneously from the soma with a pipette filled with Alexa 488 (50 μM) to visualize the axon, and from the axon cut end (i.e. bleb) at distances ranging from 50 to 152 μm (mean: 92 ± 19 μm, n = 9). In accordance with the increased Ca^2+^ influx during ISF, amplitude of si-AP was found to be increased in the axon (by 12.4 ± 0.8 mV, n = 9; Wilcoxon-test p < 0.05; **Fig. 4a**) but not in the soma (difference: 1.0 ± 1.2 mV, n = 9; Wilcoxon-test, p > 0.1). Moreover, the axonal spike amplitude followed an exponential decay as a function of pre-pulse duration that is similar to the decay of synaptic strength found in Fig. 1C (time-constant of 4.0 ms; **Supplementary Fig. 4**). However, the half-width of the axonal si-AP was not significantly broader than the ai-AP (105 ± 3%, n = 8; Wilcoxon test p>0.1; **Supplementary Fig. 4**). Interestingly, the spike threshold measured in the axon was hyperpolarized by ~4 mV during ISF (from −45.4 ± 2.0 to −49.7 ± 2.5 mV, n = 9, Wilcoxon-test p < 0.05; **Fig. 4b**) and the maximal slope of the rising phase of the spike was increased (from 206 ± 19 to 251 ± 21 mV/ms, n = 9, Wilcoxon-test p < 0.01; **Fig. 4b**), indicating that voltage-dependent sodium (Na_v_) channel inactivation is minimized when the spike is induced by synchronous inputs. We next compared ISF with another synaptic facilitation that also depends on an increase in presynaptic spike amplitude following transient hyperpolarization (h-ADF) ^17^. Compared to h-ADF, ISF was found to be more robust at both L5-L5 (125 ± 8%, n = 7 vs. 106 ± 3%, n = 7, p<0.05) and CA3-CA3 connections (126 ± 7% n = 9 vs. 104 ± 3%, n = 9, p<0.05; **Supplementary Fig. 5**). This difference probably results from the larger enhancement of the axonal spike overshoot during ISF compared to that found during h-ADF (respectively, 7.2 ± 1.2 mV, n = 8 and 5.5 ± 1.5 mV, n = 5). Altogether, these data indicate that modulation of spike amplitude is at the origin of ISF through a Ca^2+^-dependent mechanism ^19^.

**Fig. 4.**
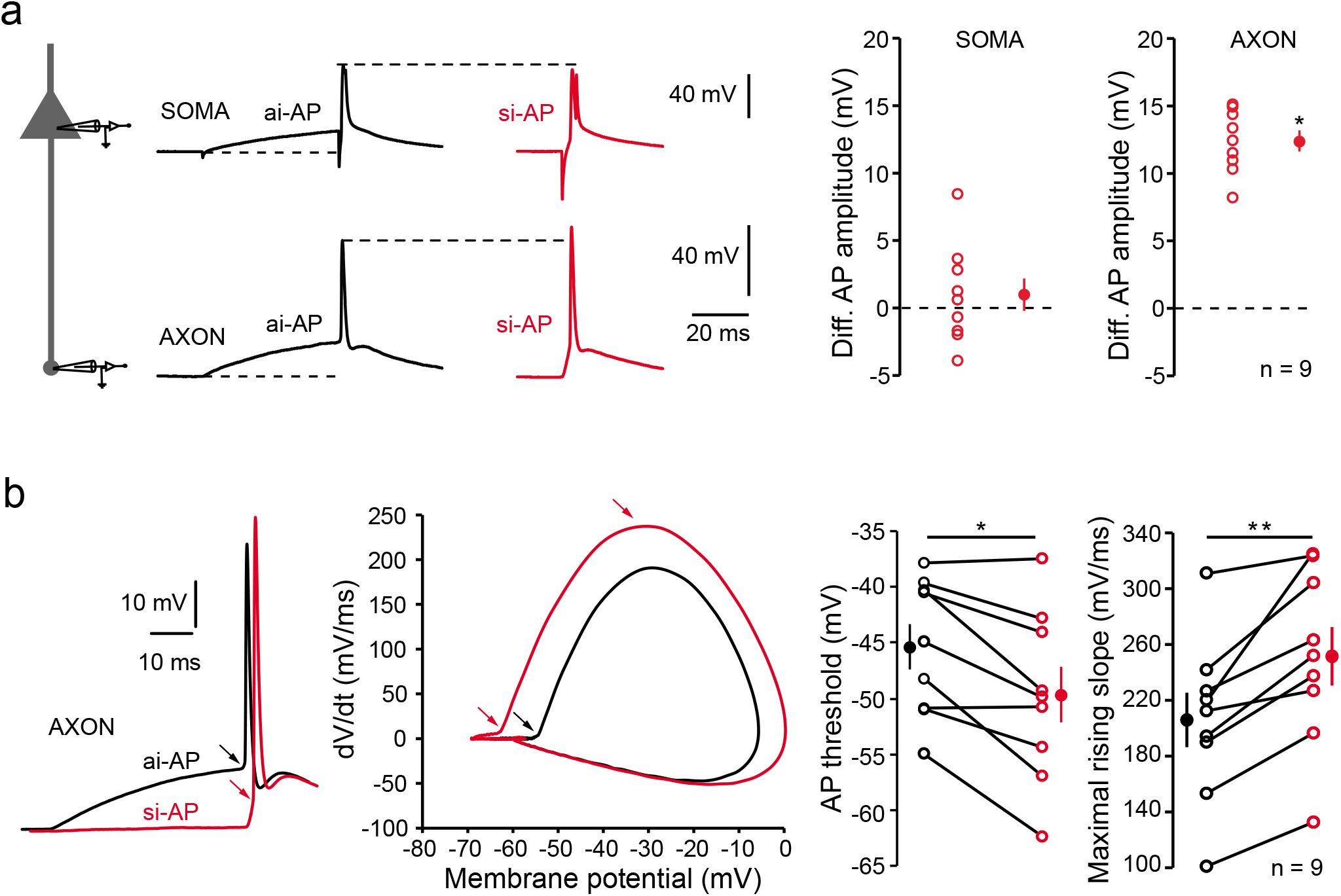
Modulation of axonal spike amplitude. **a**, ISF is associated with a modulation of axonal spike amplitude. Left, recording configuration. Simultaneous whole-cell recordings from the soma and cut end axon in L5 pyramidal cells were obtained. Middle, comparison of somatic and axonal action potential amplitudes evoked by asynchronous-like (black traces) and synchronous-like (red traces) inputs. Note the increased amplitude in the axon but not in the soma. Right, quantitative data. **b**, Hyperpolarization of spike threshold and increase in the slope of the spike rising-phase during ISF. Left, comparison of axonal spikes induced by asynchronous and synchronous inputs. Middle, phase-plot of the action potentials. Note the hyperpolarization of the threshold during and the increase of maximal slope during ISF. Right, quantitative data.

### Role of sodium channel inactivation

The differential sensitivity of the spike amplitude in the axonal and the somatic compartment to depolarization may result from the difference in Na_v_ channel inactivation in these two compartments. In fact, axonal Na_v_ channels display a hyperpolarized inactivation curve (half inactivation of −80 mV) and are highly sensitive to inactivation by subthreshold depolarization^20^. We used a NEURON model of a fully reconstructed L5 neuron ^20^ to determine the role of axonal Na_v_ channel inactivation in ISF. Synaptic conductance was injected in the soma in the form of 5 inputs whose delay varied between 0 and 10 ms to simulate different input synchrony. Presynaptic Na_v_ channel inactivation, spike-evoked Na^+^ current and AP overshoot were measured in each condition in the presynaptic terminal located at 150 μm from the soma. Increased input synchrony leads to a decrease in Na_v_ channel inactivation and therefore to larger Na^+^ current and spike overshoot (**Fig. 5a**). Next, we used the model of an L5 neuron connected to a postsynaptic cell by a synapse to determine the incidence of Na_v_ channel availability on ISF. In agreement with the experiments, a si-AP had a larger amplitude and evoked a larger postsynaptic response than an ai-AP in control conditions (i.e. when half inactivation of axonal Na_v_ channels was set to – 80 mV^20^; **Fig. 5b**). Interestingly, the difference between si-AP and ai-AP amplitude was enhanced if inactivation of axonal Na_v_ channels was increased by hyperpolarizing their basal inactivation curve (half inactivation at −85 mV; **Fig. 5b**), leading to enhanced ISF. We verified experimentally the model prediction by using carbamazepine (CBZ, 100 μM), an antiepileptic drug which hyperpolarizes Na_v_ half-inactivation by ~6 mV ^21, 22^. In L5 connections, addition of CBZ reduced synaptic transmission to 56 ± 9% of the control amplitude (n = 7, Wilcoxon-test, p < 0.05; **Fig. 5c**) and increased the magnitude of ISF by 23 ± 8% (n = 7, Wilcoxon-test, p < 0.05; **Fig. 5c**).

**Fig. 5.**
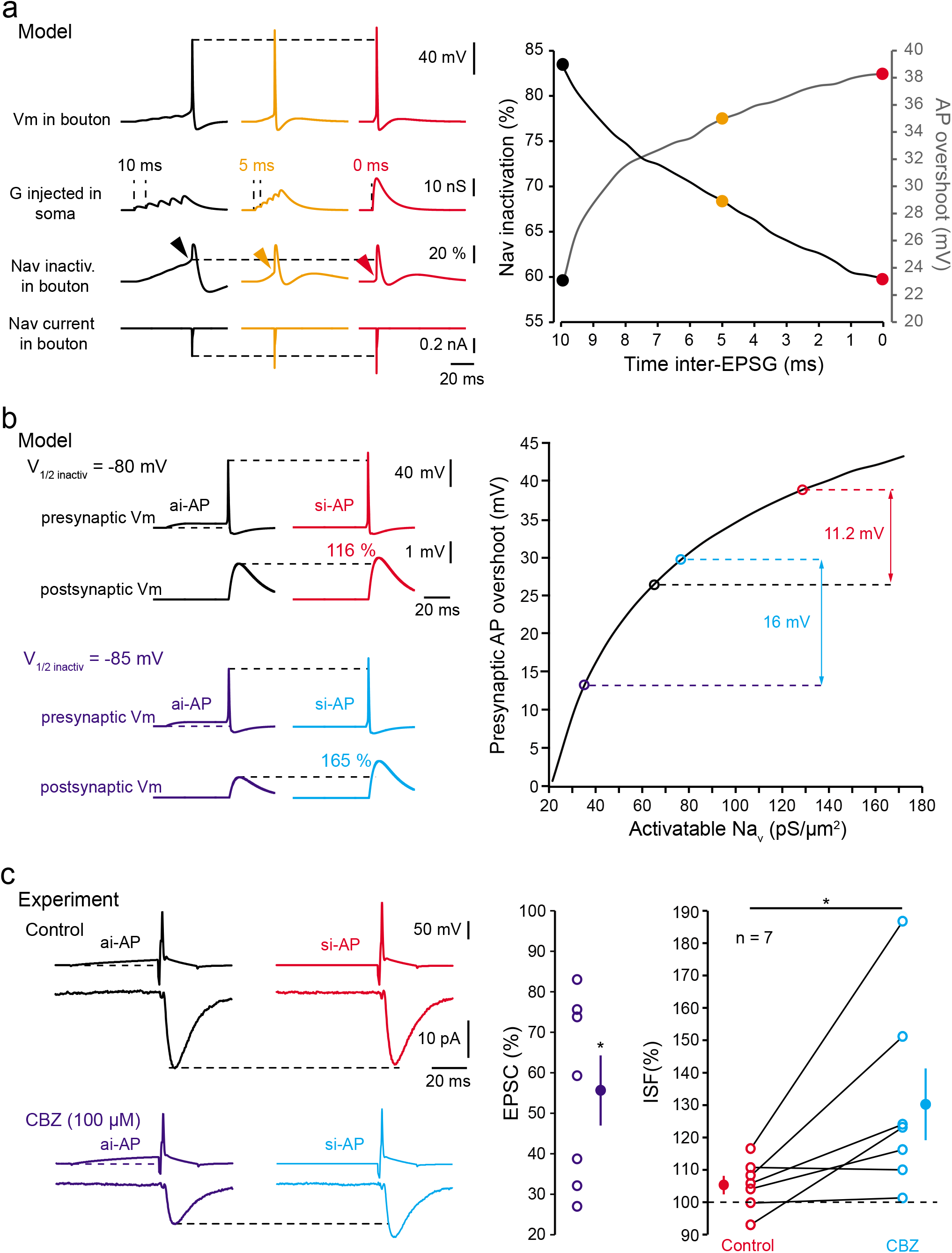
Minimization of Na_v_ channel inactivation. **a**, Reduced Na_v_ channel inactivation during ISF in a model L5 pyramidal cell. Left, voltage (top line), Na_v_ channel inactivation (middle line) and presynaptic Na^+^ current (bottom line) traces for asynchronous (black), intermediate (orange), or synchronous (red) inputs. Note the increase in spike and Na_v_ current amplitude, and the decreased Na_v_ channel inactivation measured before the spike. Right, plot of presynaptic Na_v_ channel inactivation and presynaptic spike overshoot as a function of synaptic input synchrony. **b**, Increasing presynaptic Na_v_ channel inactivation enhances ISF in a Hodgkin-Huxley model of an L5-L5 connection. Top left, control conditions (V_1/2 inacti_v = −80 mV). A presynaptic AP evoked by a synchronous input is 11.2 mV larger than that produced by an asynchronous input. This larger presynaptic spike produces a larger EPSP (ISF = 116%). Bottom left, test conditions (V_1/2 inactiv_ = −85 mV). Because Na_v_ channel inactivation is larger, the modulation of presynaptic spike amplitude (16 mV) as well as ISF (165%) is enhanced. Right, plot of the amplitude of presynaptic spike overshoot as a function of Na_v_ channel availability. **c**, Experimental verification of the role of Na_v_ channel inactivation during ISF. Top left, example of L5 connection showing no ISF in control conditions. Bottom left, same connection in the presence of carbamazepin (CBZ). The evoked EPSC is ~40% smaller and ISF is now clearly visible. Middle, plot of EPSC reduction caused by CBZ. Right, plot of ISF in control (red) and in the presence of CBZ (light blue).

To confirm the effect of changing Na_v_ channel availability on ISF in the model, we next reduced the density of Na_v_ channels in the axon from 300 pS/μm^2^ to 150 pS/μm^2^ while keeping all other parameters constant. In this condition, ISF was enhanced from 116 to 165% (**Supplementary Fig. 6**). We tested experimentally the second prediction of the model by partially blocking Na_v_ channels with a low concentration of tetrodotoxin (TTX; 20-40 nM). TTX reduced synaptic transmission to 75 ± 9% of the control value (n = 7; **Supplementary Fig. 6**) and increased ISF by 10 ± 2% (n = 7; Wilcoxon-test p < 0.05). We conclude that ISF results from the minimization of Na_v_ channel inactivation by synchronous inputs that in turn leads to the increase in presynaptic AP amplitude, calcium influx, and glutamate release.

### ISF induced in a single neuron affects network activity

Next, we tested whether ISF triggered in a single neuron had a significant impact on spontaneous activity in cortical circuits. For this, we approached the physiological conditions of extracellular calcium (i.e. 1.3 mM external calcium instead of 3 mM ^23, 24^). First, we measured the effect of reducing external calcium on ISF. We found that decreasing extracellular calcium concentration from 3 mM to 1.3 mM decreased synaptic strength (to 40 ± 8% of the control EPSC, n = 8; Wilcoxon-test, p<0.01) but increased ISF amplitude (from 106 ± 4% to 118 ± 6%, n = 8; Wilcoxon-test, p < 0.01; **Supplementary Fig. 7**). In a second step, paired-recordings from adjacent L5 pyramidal neurons were obtained and a spike was elicited in one of the two cells. The spontaneous activity was measured in the non-spiking cell by counting the number of post-synaptic events occurring 200 ms before the spike and 200 ms after the spike (**Fig. 6a**). In order to verify the integrity of the axon, neurons were filled with Alexa 488 and recordings from cells with damaged axons were rejected. We found that spontaneous activity in the non-spiking cell increased after a si-AP but not after an ai-AP (132 ± 7% versus 99 ± 4%, n = 9; Wilcoxon-test p < 0.01; **Fig. 6b**). We conclude that ISF induced in a single neuron significantly affects spontaneous activity in local cortical circuits suggesting that transmission of input synchrony through ISF would represent a robust way for information coding.

**Fig. 6.**
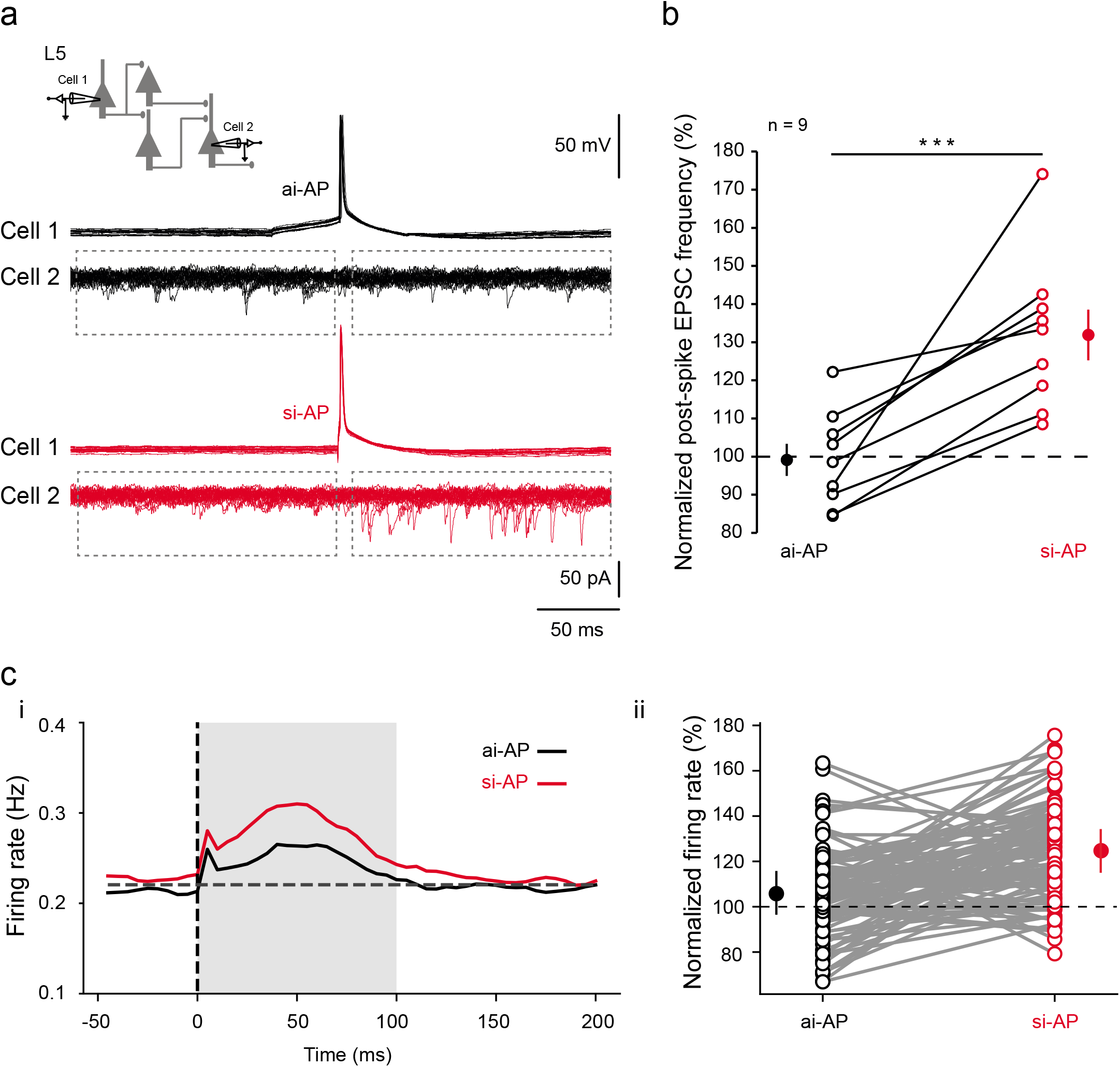
Effects of ISF on network activity. **a**, Incidence of ISF on cortical network activity. Spontaneous activity was measured in cell 2 before and after the spike triggered in cell 1 and the event frequency post-spike was normalized to that measured pre-spike. Black traces, no change in network activity is induced by ai-APs (99%). Red traces, increased network activity induced by si-APs (124%). **b**, Plot of normalized EPSC frequency triggered by ai-APs (black circles) and by si-APs (red circles). **c**. Effect of ISF in a balanced network model. **i.** Averaged firing rate (over 100 repeats on 100 neurons and over 5 networks) just before and after the injection of an ai-AP spike (black curve) and a si-AP spike (red curve). **ii.** Individual normalized changes in spiking activity, for the 100 neurons in a given network, when injecting an ai-AP (black) and a si-AP (red).

To confirm this result, we used a computational approach. We built a network model made of 4000 excitatory cells and 1000 inhibitory cells randomly connected with a probability of 10% (**Supplementary Fig. 8**). Neurons present a low spiking activity around 0.2 Hz. The network was balanced for excitatory and inhibitory synaptic weights (see Methods for details). We reproduced the protocol used in the experiments to check the effect of ISF on the network. One excitatory neuron was randomly chosen and was forced to fire one extra spike. To model a si-AP, EPSPs induced by this extra spike were increased by 30% in target neurons. For an ai-AP, EPSPs were left unchanged. The spiking rate of the network was measured 100 ms before and 100 ms after the extraspike. This protocol was repeated for 100 ai-APs and 100 si-APs for a given neuron, and results over 5 random networks were averaged. Overall, we found that an ai-AP produced on average an increase of 8.7% of the network spiking activity while a si-AP induced on average an increase of 25.2% (**Fig. 6c**). This result confirms our experimental observation that ISF induced in a single neuron can affect local circuit activity.

### Discussion

We show here that sodium channels read and transmit levels of input synchrony with a high temporal precision to local cortical circuits through the modulation of presynaptic action-potential amplitude. Synaptic transmission was found to be facilitated at L5-L5 or at CA3-CA3 connections when the presynaptic action potential was triggered by highly synchronous inputs or by a short pulse of current that produced in all cases a steep depolarization from resting membrane potential. Using dual soma-axon recordings, calcium imaging, pharmacological tools and computer modeling, we demonstrate that this facilitation results from an elevation in presynaptic calcium due to the enhancement of axonal spike amplitude caused by minimized inactivation of axonal Na_v_ channels. This input synchrony-dependent facilitation (ISF) was found to enhance local circuit activity, suggesting that it may represent a robust way of transmitting synchrony in the brain.

So far, spike-threshold adaptation reported in cortical neurons was thought to represent the most accurate mechanism for detection of fast input synchrony in central neurons ^7–10^. We suggest here that it may represent only the emerged tip of the iceberg because spike-threshold dynamics appears to be only an indicator of Nav channel inactivation in the presynaptic cell and the consequences on the network can only be evaluated by paired intracellular recording from connected neurons, a particular challenging technique *in vivo*. In fact, the most important consequence of input synchrony-dependent adaptation in spike-threshold is an enhancement of presynaptic release caused by modulation of presynaptic spike amplitude. As a consequence, input synchrony both increases cell excitability by lowering the spike threshold and synaptic transmission via the enhancement of the spike amplitude in the axon.

The contribution of axonal Na_v_ channels in input-synchrony-dependent modulation of presynaptic release is shown by four lines of evidence. First, whole-cell recording from the axon showed that the spike amplitude in the axonal compartment was highly modulated. In particular the maximal slope of the rising phase of the spike was increased. Second, computer simulation showed that Na_v_ channel inactivation caused by subthreshold depolarization preceding the spike decreased substantially when input synchrony increased. Third, enhancement of Na_v_ channel inactivation by CBZ was found to enhance the magnitude of ISF. Finally, reduction of Na_v_ channel availability by TTX increased ISF. Importantly, the implication of Na_v_ channels had been shown in another context-dependent enhancement of synaptic transmission that depends on presynaptic hyperpolarization ^17^. This facilitation was due to recovery of axonal sodium channels from inactivation. Therefore, availability of presynaptic Na_v_ channels represents a major determinant of synaptic transmission at local excitatory connections.

The modulation of release we report here is due to a Na_v_ channel-dependent enhancement of presynaptic action potential amplitude and calcium influx. The fact that increasing presynaptic spike amplitude enhances calcium entry and neurotransmitter release has been shown in many axon types. For example, in cultured hippocampal neurons, voltage-imaging of presynaptic boutons indicates that action potentials display a very small overshoot ^25^, a condition favorable for the modulation of Ca^2+^ entry by changes in AP amplitude. Moreover, voltage-clamp recordings from presynaptic boutons in cerebellar cultured neurons show that enhanced spike amplitude results in larger presynaptic Ca^2+^ currents and in enhanced release ^19^. In addition, partial blockade of Na_v_ channels with low concentration of TTX both reduces AP amplitude and synaptic release in connected pairs of L5 pyramidal neurons ^26^.

ISF enters into the group of context-dependent modulations of synaptic release that depends on axonal ion channels. To date, 3 types of modulation have been identified: 1) a K_v_1 channel-dependent form of synaptic facilitation induced by long presynaptic depolarizations preceding the presynaptic AP ^14, 15, 27, 28^, 2) a Ca_v_2.1 channel-dependent accumulation of presynaptic Ca^2+^ induced by slow presynaptic depolarizations that facilitates presynaptic release in short cerebellar axons ^29, 30^ and 3) a Na_v_-dependent facilitation induced by presynaptic hyperpolarization ^17^. Among these context-dependent modulations of synaptic transmission, only the last one may have a significant impact on the activity of fast cortical circuits, the two other processes being too slow and probably involved in slower network activity ^14, 15, 27, 28^. While both ISF and Kv1-dependent modulation of synaptic release have been observed on the same synapses, their respective time-windows are clearly distinct (in the millisecond range for ISF and in the second range for Kv1-dependent modulation). The comparison of hyperpolarization-induced facilitation and ISF on the same synapses show larger effects produced by ISF, probably due to a larger enhancement of the spike overshoot.

From a functional point of view, ISF can be seen as a time coding process in which action potentials elicited with a short latency would produce stronger responses than those induced with a long latency. Theoretical work suggests that coding based on absolute spike time is by far the most powerful because of its high capacity for information transfer compared to rate coding or rank coding ^31^. In addition, several experimental studies indicate that the precise timing of the first spike carries critical information about the stimulus in sensory responses ^32–35^. Our data indicate that ISF conveys presynaptic spike timing information to the postsynaptic cell with high temporal precision and thus provides strong biological support for time coding based on spike latency at cortical synapses ^31, 36^. ISF defines a rule in which highly relevant neuronal information comes early and is transmitted in the circuit while poorly relevant information comes later and is filtered. This rule is reminiscent of the principle governing the release-dependent variation in synaptic delay at unitary L5-L5 synapses where large synaptic responses display the shortest synaptic delays and small responses, the longest ones ^37^.

Does ISF also occur *in vivo*? Although a direct answer to this question is technically extremely challenging, we assume that this might be the case. In fact, visually evoked inputs differentially inactivate Na_v_ channels and therefore highly modulate spike-threshold in primary visual cortical neurons ^7^. In the barrel cortex, preferred direction elicits steeper EPSPs that in turn evoke spikes with a lower threshold ^11^. Thus, spikes evoked by highly synchronous inputs (or fast rates of depolarization) are likely to better transmit information in the network than spikes evoked by weakly synchronous inputs (or slow rates of depolarization). Our experimental results show that this might be the case. Indeed, spikes elicited by synchronous-like inputs have a stronger impact on network activity compared to spikes elicited by asynchronous-like inputs.

The magnitude of ISF reported in the present study is relatively modest (20-40%). There are, however, good reasons to believe that ISF could be significantly larger *in vivo*. First, we show that ISF is enhanced in physiological extracellular Ca^2+^. Second, the presence of neuromodulators such as acetylcholine would significantly promote ISF. In fact, acetylcholine specifically depolarizes axon membrane potential by blocking Kv7 channels ^38^ and reduces sodium current ^39^, which should increase ISF (see TTX experiment, Suppl. Fig 6). But the contribution of ISF to network dynamics might be even larger during early development. Indeed, the low density of Na_v_ channels together with the slow kinetics of EPSPs observed during development represent two factors that may optimize ISF. Further experimental investigations will be needed to determine the importance of ISF in the cortex *in vivo*.

## Acknowledgments

Supported by INSERM, CNRS (DD), Ecole Normale Supérieure (doctoral grant to MZ), Agence Nationale de la Recherche (grant ANR-14-CE13-0003 to DD, RB and PY) and Fondation pour la Recherche Médicale (DVS-2013-1228768 to DD and FDT-2015-0532147 to MZ). We thank A Bialowas & V Marra for sharing preliminary data, Urs Gerber, Gilles Laurent, Erwin Neher & Wolf Singer for critically reading a preliminary version of the manuscript and Rosa Cossart & Massimo Scanziani for helpful discussion.

## Author contributions

MZ, SR & DD conceived the project, MZ, SR, YI, NBG, LFM & MR performed the experiments, MZ, SR, YI & DD analyzed the data, MZ, PY & RB performed modelling and MZ, SR, PY & DD wrote the manuscript.

## Additional information

Supplementary Fig.1-8.

Supplementary Table 1

---

**Supplementary Fig. 1.**
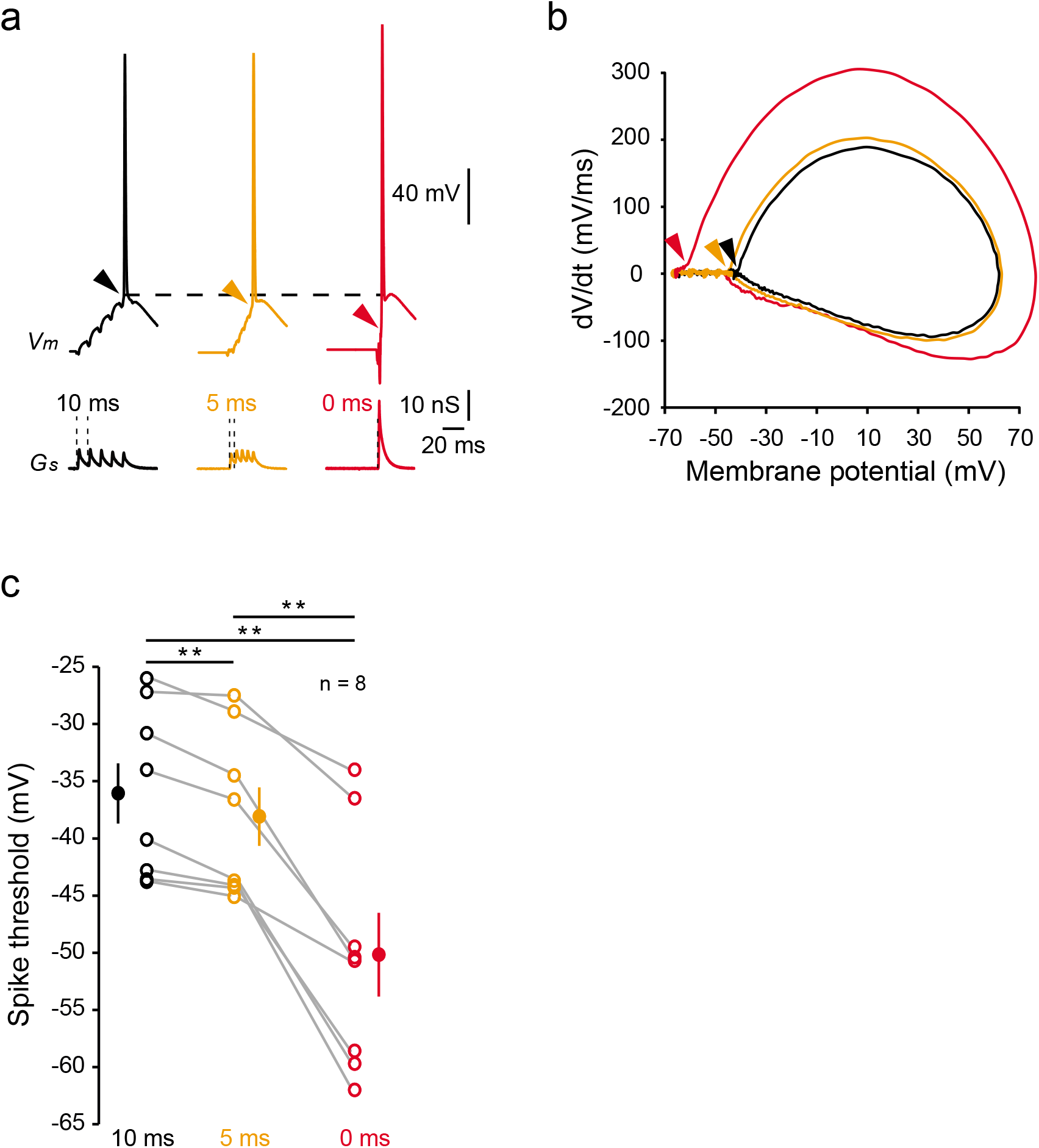
Input synchrony lowers presynaptic AP threshold. **a**, Experimental paradigm and action potentials triggered by various degree of input synchrony (black: 10 ms, orange: 5 ms and red: 0 ms). **b**, Phase plots of the APs in the 3 conditions, note the hyperpolarization of the AP threshold (arrow head) for the maximal input synchrony (red). **c**, Data analysis. **, Wilcoxon test, p<0.01.

**Supplementary Fig. 2.**
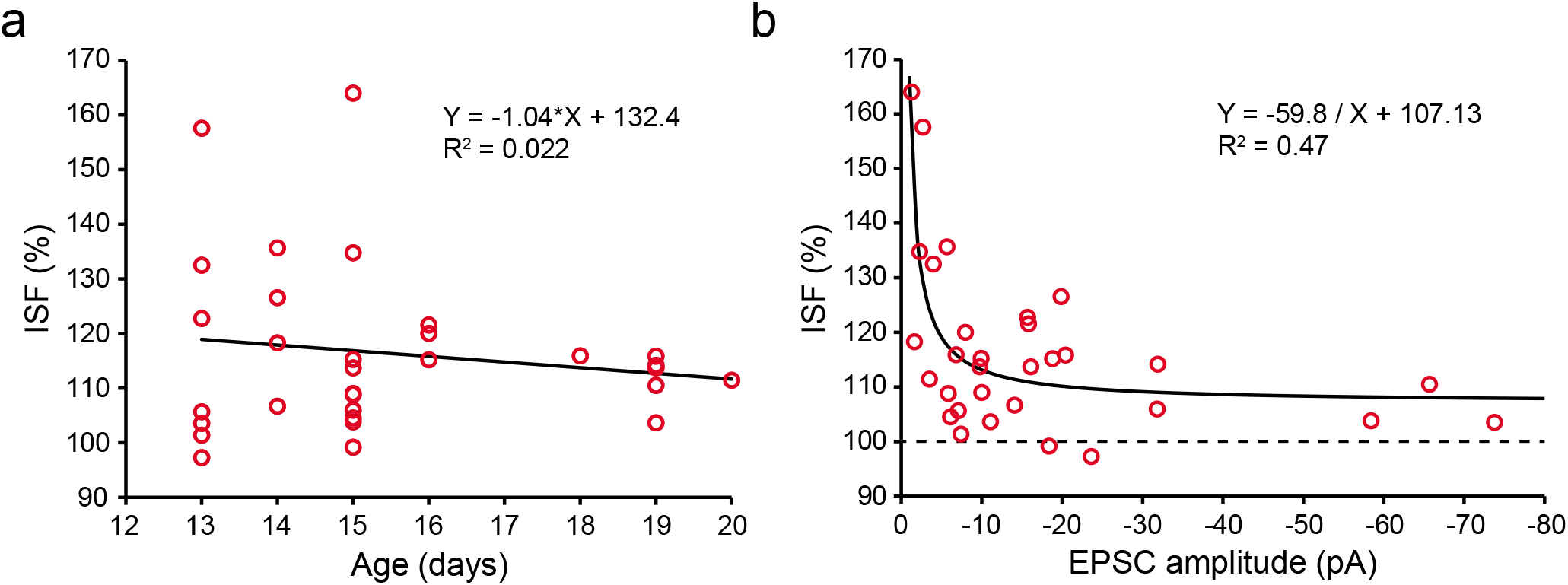
Properties of ISF. **a** & **b**, Age- and amplitude-dependence of ISF. Left, plot of ISF as a function of age. Black line, linear regression (y = −1.04*x + 132.42, R^2^ = 0.022). Right, plot of ISF as a function of EPSC amplitude. Black curve, regression (y = −70.37/x + 107.67, R^2^ = 0.322).

**Supplementary Fig. 3.**
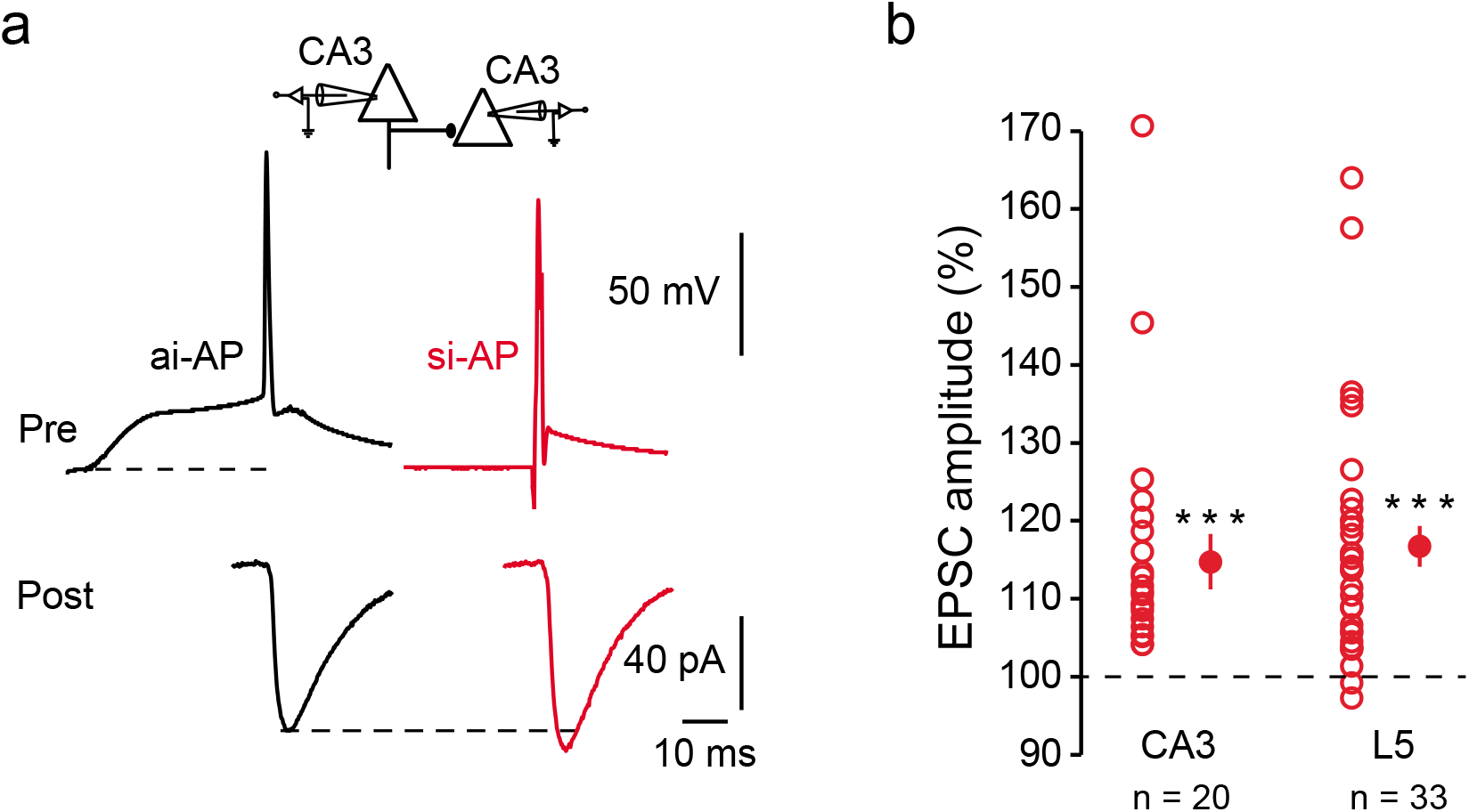
ISF at CA3-CA3 synapses. **a**, Representative example of electrophysiological recordings from a connected pair of CA3 neurons. Note the increase in postsynaptic response amplitude evoked by an action potential produced by synchronous-like input (red). **b**, Comparison of ISF in CA3 (n = 20) and L5 (n = 33) neurons (plots of EPSC amplitude evoked by synchronous-like inputs have been normalized to those evoked by asynchronous-like inputs). ***: Wilcoxon test, p<0.001.

**Supplementary Fig. 4.**
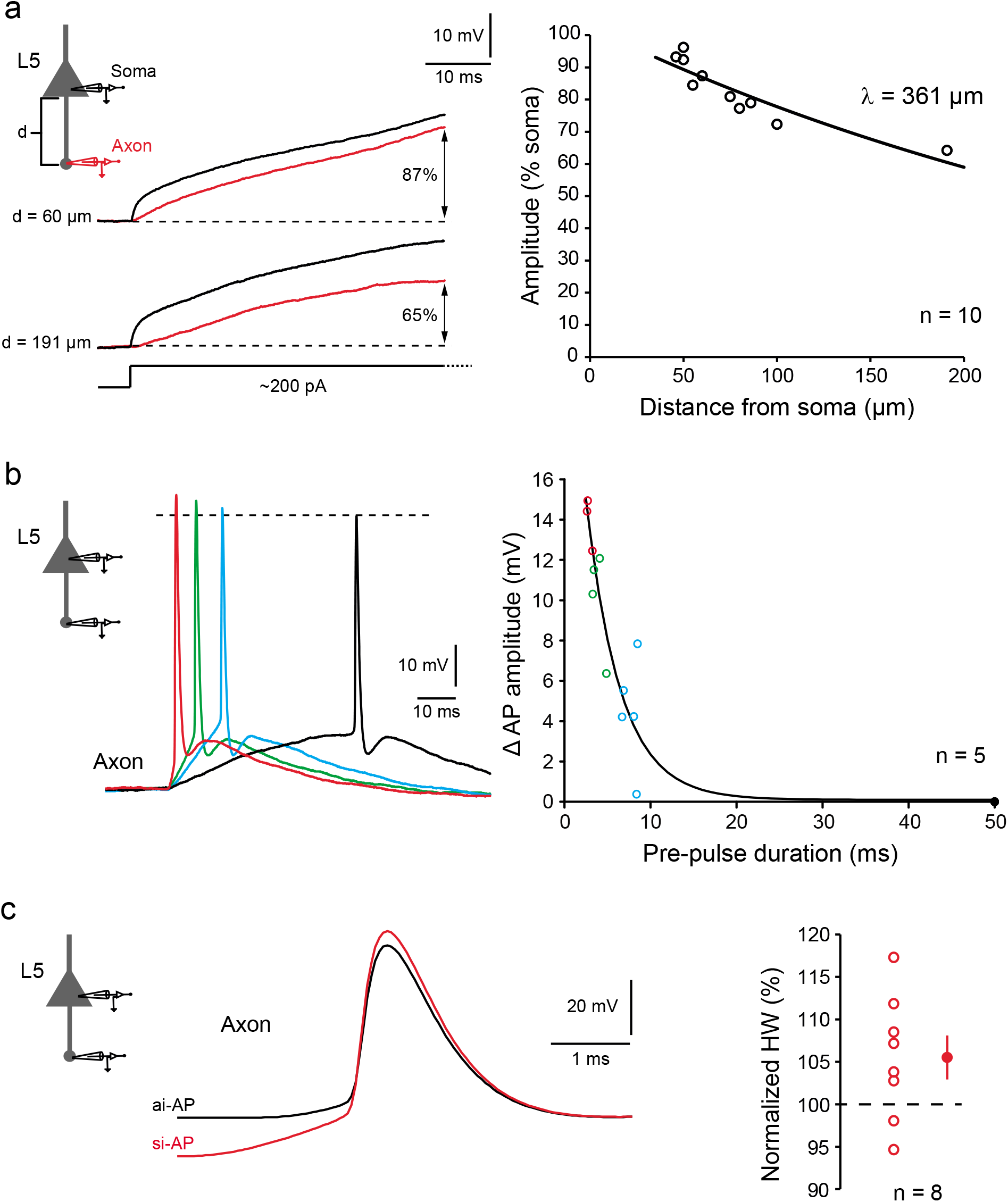
Soma-axon recording: spatial and temporal properties. **a**, Attenuation of subthreshold depolarization along the axon. Left, voltage traces recorded simultaneously in the soma (black) and the axon (red) upon current injection in the cell body. Note the larger attenuation observed for the longest axonal recording distance. Right panel, exponential fit of the depolarization measured in the axon normalized to that measured in the soma (*y* = 102.6 ∗ *e^−x/λ^*, with λ = 361 μm, R^2^ = 0.83). **b**, Time-course of enhanced spike amplitude recorded in the axon. Left, action potential evoked by asynchronous-like (black), intermediate-like (blue & green), and synchronous-like (red) inputs. Right, variation in spike amplitude as a function of spike latency (5 axons). Each data point of the same color represents averaged values in a single recording. Note the rapid modulation in spike amplitude (black line, exponential regression (*y* = −0.098 + 28.12 ∗ *e^−x/τ^*, with τ = 4.0 ms, R^2^ = 0.91). **c**, No change in axonal spike duration was observed during ISF (p>0.1). Left, representative example (black trace: ai-AP, red trace: si-AP). Right, pooled data.

**Supplementary Fig. 5.**
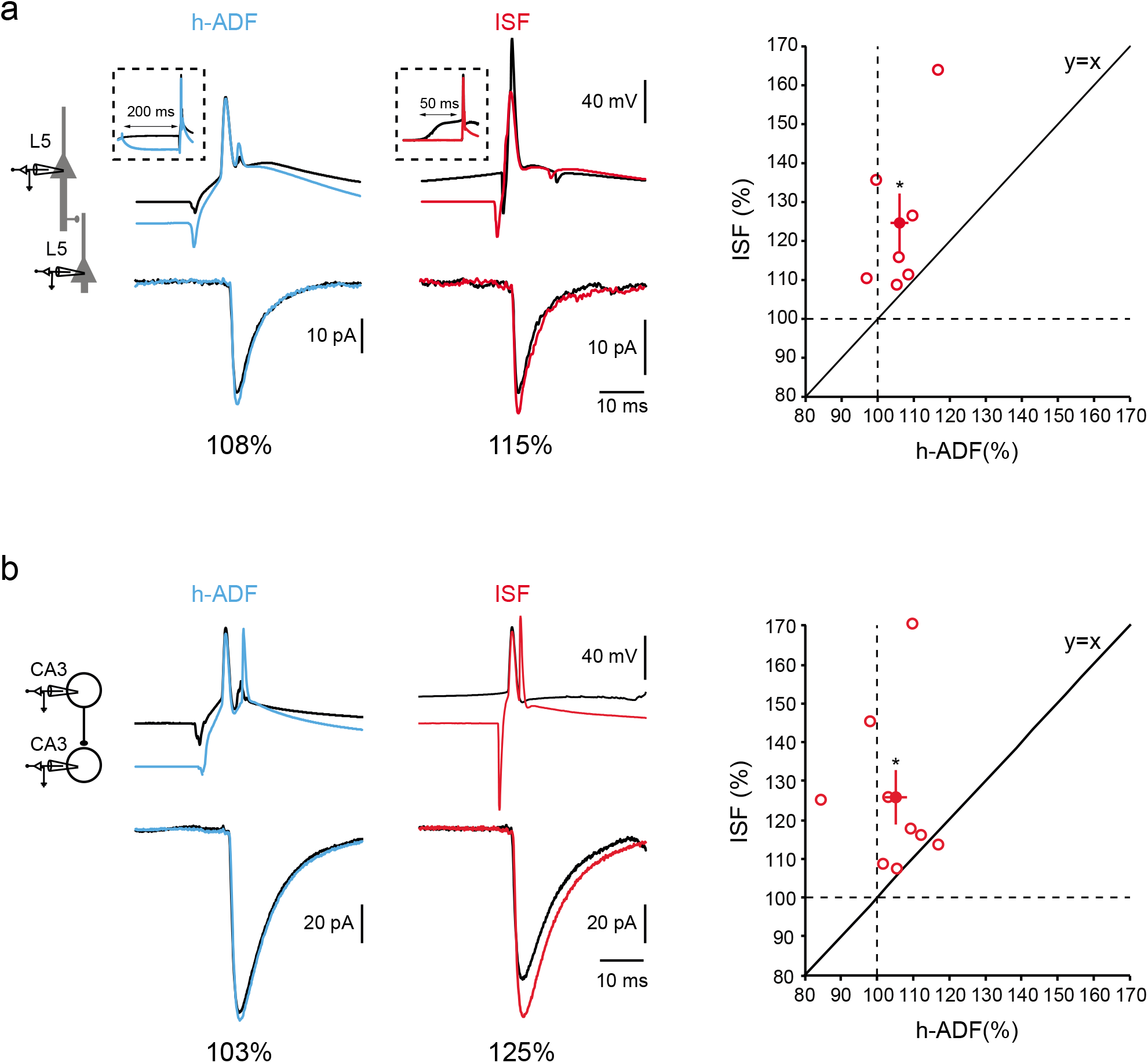
Comparison of ISF and h-ADF. **a**, L5-L5 synapses. Left, h-ADF induced by transiently hyperpolarizing the membrane potential amounts to 108% of the control EPSC. Right, in the same pair of neurons ISF amounts to 115%. Right, pooled data over 8 L5-L5 cell pairs showing greater facilitation for ISF compared to h-ADF. **b**, CA3-CA3 synapses. Left, representative example showing a small h-ADF (103%) for a large ISF (125%). Right, pooled data over 9 CA3-CA3 cell pairs showing greater ISF compared to h-ADF.

**Supplementary Fig. 6.**
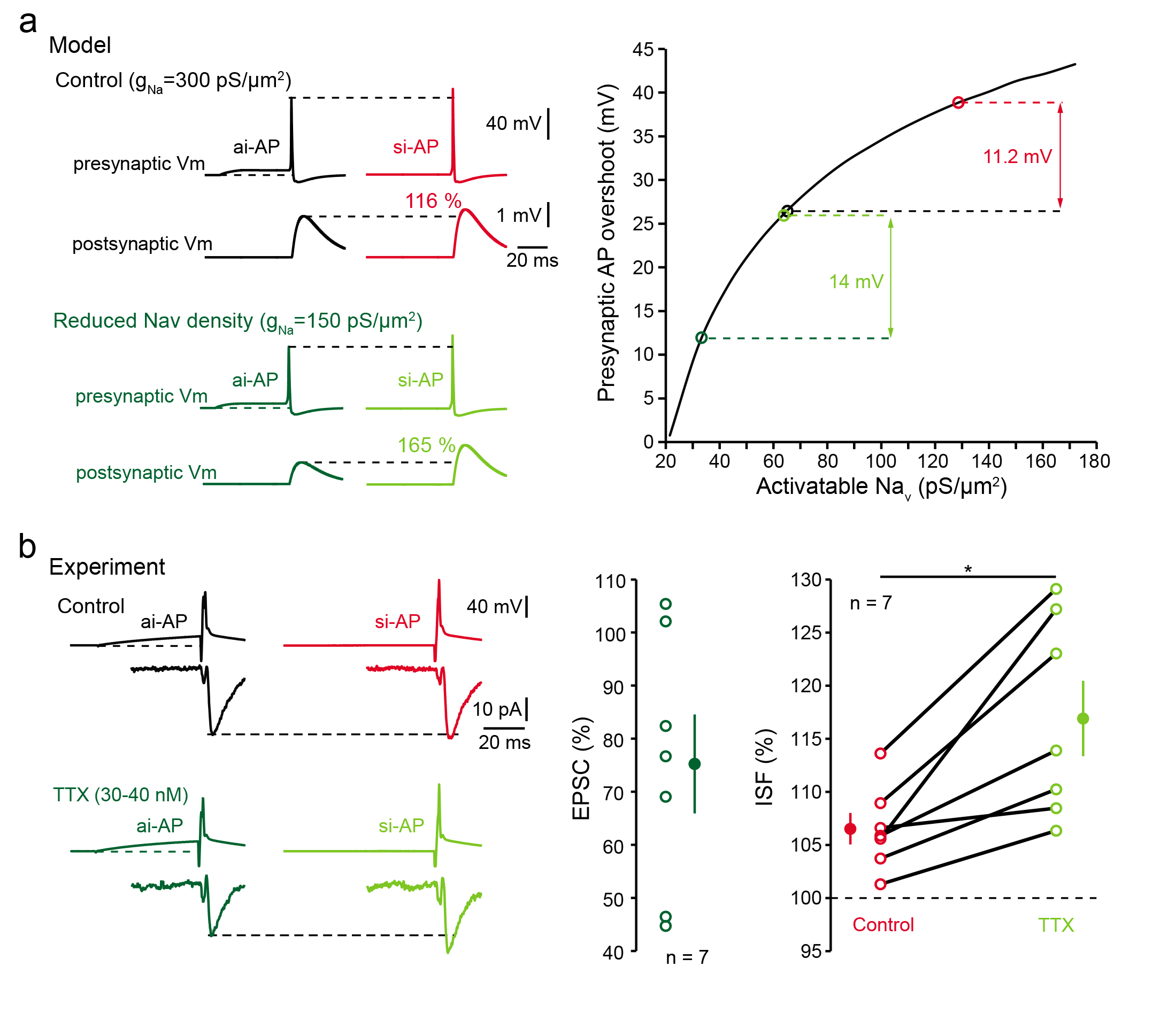
Modulation of ISF by Na_v_ density. **a**, Modeled reduction of Na_v_ density and enhancement of ISF. Top left, control conditions (g_Nav_ = 300 pS/μm^2^). Bottom left, the reduction of Na_v_ density by 50% (g_Nav_ = 150 pS/μm^2^) enhances the modulation of presynaptic spike amplitude and ISF. Right, presynaptic spike amplitude as a function of activatable Na_v_ channel density. Note that the non-linear relationship accounts for increased modulation of spike amplitude during ISF (red in control condition and light green in low Na_v_ channel density condition). **b**, Experimental verification of theoretical prediction. Top left, ISF in control condition. Bottom left, enhancement of ISF in the presence of tetrodotoxin (TTX). Middle, plot of EPSC amplitude reduction caused by TTX. Right, pooled data showing the increase of ISF in TTX.

**Supplementary Fig. 7.**
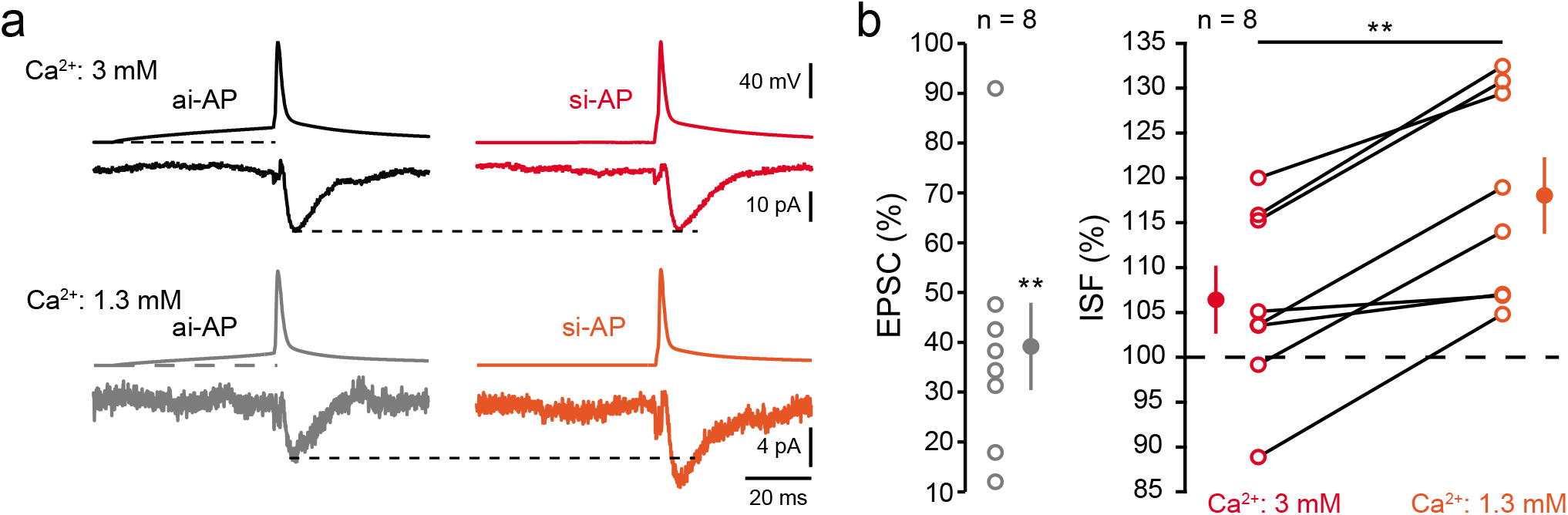
ISF in physiological external concentration of Ca^2+^. **a**, Top, lack of ISF in 3 mM Ca^2+^. Bottom, ISF in 1.3 mM Ca^2+^ (same connection). **b**, Left, plot of EPSC amplitude in 1.3 mM Ca^2+^ normalized to the control situation (3 mM Ca^2+^). Right, pooled data of ISF in 3 mM Ca^2+^ and 1.3 mM Ca^2+^.

**Supplementary Fig 8.**
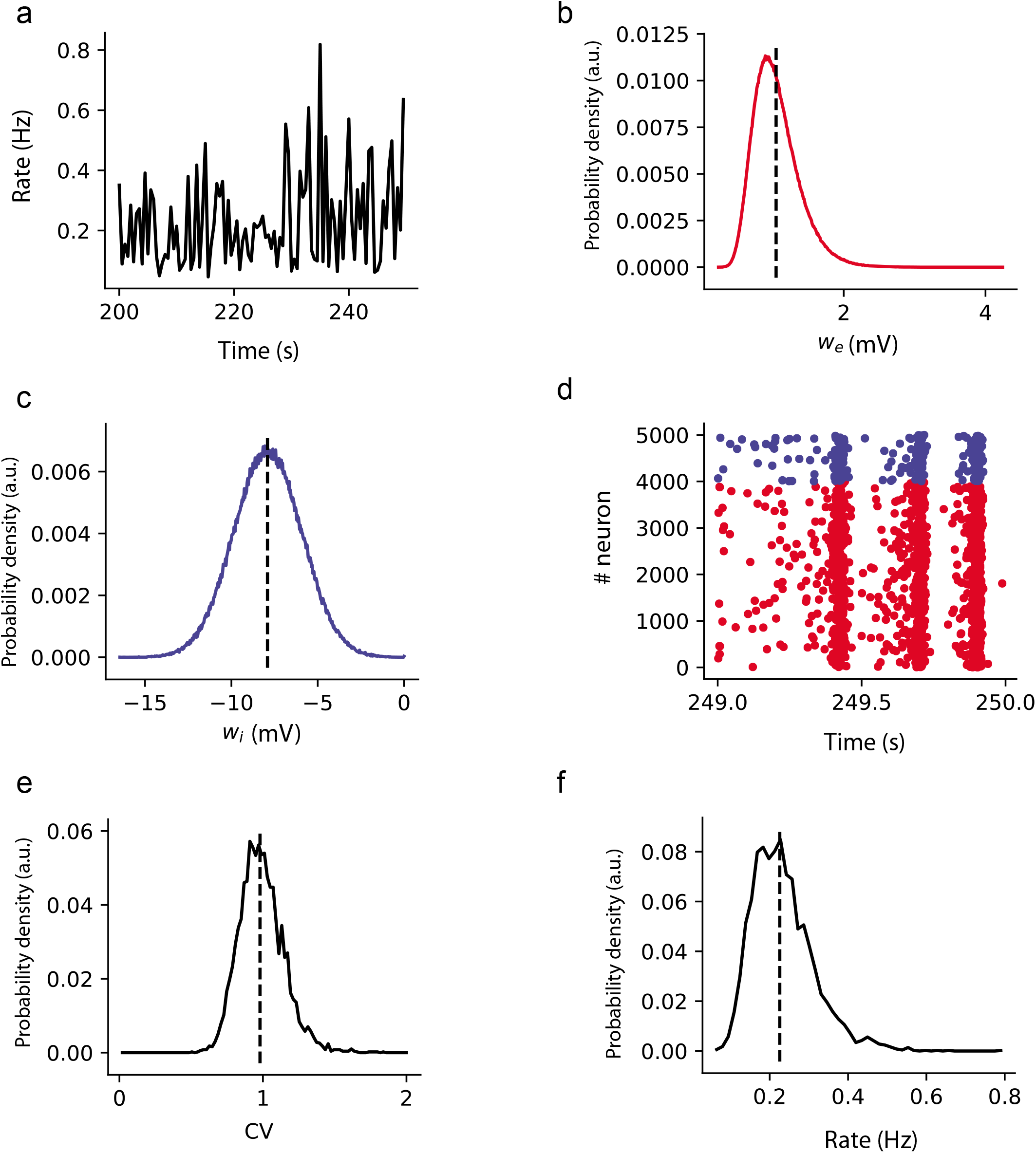
Dynamics of the balanced network. **a.** Firing rate of the whole network during 50 s. **b.** Distribution of the excitatory weights. Dash dotted line shows the mean weight < *w_exc_* > = 1 *mV* **c.** Distribution of the inhibitory synaptic weights. Dash-dotted line shows the mean weight. **d.** Raster plot of the activity during 1 second for all neurons (excitatory in red, inhibitory in blue). **e.** Distribution of the coefficients of variation for the inter-spike intervals, over all neurons in the network. Dash-dotted line shows the mean CV, equal to 1. **f.** Distribution of the firing rates over all neurons in the network. Dash dotted line shows the mean at 0.21Hz.

**Supplementary Table 1.**
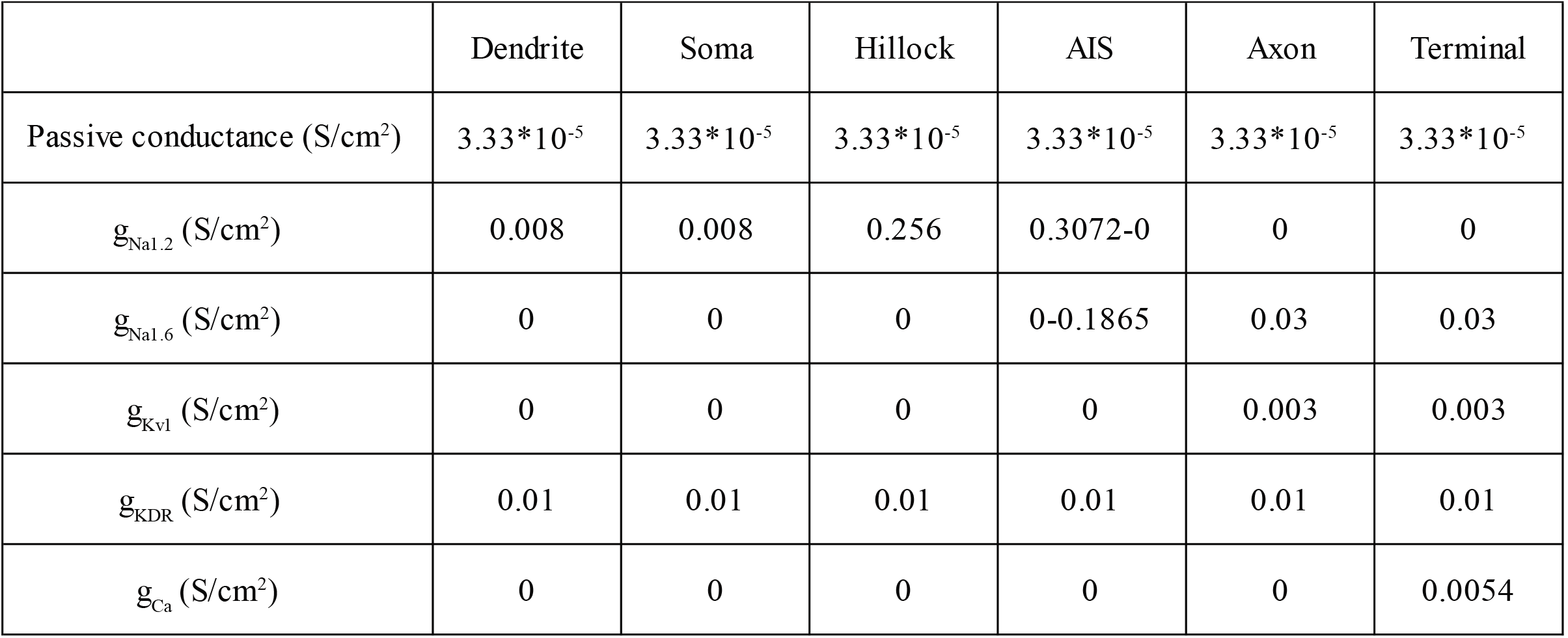

|  | Dendrite | Soma | Hillock | AIS | Axon | Terminal |
| --- | --- | --- | --- | --- | --- | --- |
| Passive conductance (S/cm<sup>2</sup>) | 3.33*10<sup>-5</sup> | 3.33*10<sup>-5</sup> | 3.33*10<sup>-5</sup> | 3.33*10<sup>-5</sup> | 3.33*10<sup>-5</sup> | 3.33*10<sup>-5</sup> |
| $g_{Na1.2}$ (S/cm<sup>2</sup>) | 0.008 | 0.008 | 0.256 | 0.3072-0 | 0 | 0 |
| $g_{Na1.6}$ (S/cm<sup>2</sup>) | 0 | 0 | 0 | 0-0.1865 | 0.03 | 0.03 |
| $g_{Kvl}$ (S/cm<sup>2</sup>) | 0 | 0 | 0 | 0 | 0.003 | 0.003 |
| $g_{KDR}$ (S/cm<sup>2</sup>) | 0.01 | 0.01 | 0.01 | 0.01 | 0.01 | 0.01 |
| $g_{Ca}$ (S/cm<sup>2</sup>) | 0 | 0 | 0 | 0 | 0 | 0.0054 |

## Online Methods

### Acute slices of rat neocortex

Neocortical slices (350-400 μm) were obtained from 14- to 20-day-old Wistar rats of both sexes, according to the European and Institutional guidelines (Council Directive 86/609/EEC and French National Research Council and approved by the local health authority (Préfecture des Bouches-du-Rhône, Marseille)). Rats were deeply anesthetized with chloral hydrate (intraperitoneal, 200 mg/kg) and killed by decapitation. Slices were cut in an ice-cold solution containing (mM): 92 *n*-methyl-D-glutamine (NMDG), 30 NaHCO_3_, 25 D-glucose, 10 MgCl_2_, 2.5 KCl, 0.5 CaCl_2_, 1.2 NaH_2_PO_4_, 20 HEPES, 5 sodium ascorbate, 2 thiourea and 3 sodium pyruvate, and were bubbled with 95% O_2_-5% CO_2_, pH 7.4. Slices recovered (1 h) in a solution containing: 125 NaCl, 26 NaHCO_3_, 2 CaCl_2_, 2.5 KCl, 2 MgCl_2_, 0.8 NaH_2_PO_4_ and 10 D-glucose, and were equilibrated with 95% O_2_-5% CO_2_. Each slice was transferred to a submerged chamber mounted on an upright microscope (Olympus BX51WI or Zeiss Axio-Examiner Z1) and neurons were visualized using differential interference contrast infrared videomicroscopy.

### Organotypic slices of rat hippocampus

Hippocampal slice cultures were prepared using an interface technique^40^. Briefly, postnatal day 5-7 Wistar rats were deeply anesthetized by intraperitoneal injection of chloral hydrate, the brain was quickly removed, and each hippocampus was individually dissected. Hippocampal slices (350 μm) were placed on 20 mm latex membranes (Millicell) inserted into 35 mm Petri dishes containing 1 ml of culture medium and maintained for up to 21 days in an incubator at 34°C, 95% O_2_-5% CO_2_. The culture medium contained (in ml) 25 MEM, 12.5 HBSS, 12.5 horse serum, 0.5 penicillin/streptomycin, 0.8 glucose (1 M), 0.1 ascorbic acid (1 mg/ml), 0.4 HEPES (1 M), 0.5 B27 and 8.95 sterile H_2_O. To avoid glial proliferation, 5 μM Ara-C was added to the medium at 3 days *in vitro* (DIV) for one night. Pyramidal cells from the CA3 area were recorded at the age of 8-12 DIV.

### Paired-recordings and analysis

Dual recording from pairs of L5 neurons or CA3 neurons were obtained as previously described ^40^. The external saline contained (in mM): 125 NaCl, 26 NaHCO_3_, 3 CaCl_2_ (1.3 in specific cases), 2.5 KCl, 2 MgCl_2_ (1 in specific cases), 0.8 NaH_2_PO_4_ and 10 D-glucose, and was equilibrated with 95% O_2_-5% CO_2_. Patch pipettes (5-10 MΩ) were pulled from borosilicate glass and filled with an intracellular solution containing (in mM): 120 K gluconate, 20 KCl, 10 HEPES, 0.5 EGTA, 2 MgCl_2_, 2 Na_2_ATP and 0.3 NaGTP (pH=7.4). Recordings were performed with 2 Axopatch-2B (Axon Instruments, Molecular Devices) or a MultiClamp-700B (Molecular Devices) at 30°C in a temperature-controlled recording chamber (Luigs & Neumann, Ratingen, Germany). Usually, the presynaptic neuron was recorded in current-clamp and the postsynaptic cell in voltage-clamp. The membrane potential was corrected for the liquid junction potential (−13 mV). Both pre- and postsynaptic cells were held at their resting membrane potential (approximatively −77 mV). Presynaptic action potentials were generated by injecting brief (3 nA for 2 ms) depolarizing pulses of current at a frequency of 0.1 Hz. For synchronous input-generated action potentials (si-APs), the depolarizing pulse was evoked directly from the resting membrane potential. For asynchronous input-generated action potentials (ai-APs), the depolarizing pulse was evoked after a 50 ms-depolarization of the presynaptic cell to ~-60/-55 mV. Paired-pulse ratio (PPR) was assessed with two presynaptic stimulations (50 ms interval). Voltage and current signals were low-pass filtered (3 kHz), and sequences (200-500 ms) were acquired at 10-20 kHz with pClamp 10 (Axon Instruments, Molecular Devices). Electrophysiological signals were analyzed with ClampFit (Axon Instruments). Postsynaptic responses could be averaged following alignment of the presynaptic APs using automatic peak detection. The presence or absence of a synaptic connection was determined on the basis of averages of 30 individual traces. Pooled data are represented as mean ± se in all figures, and we used the Mann-Whitney U-test or Wilcoxon rank-signed test for statistical comparison.

### Calcium imaging

L5 pyramidal neurons were imaged with a LSM-710 Zeiss confocal microscope. For imaging calcium in the axon of L5 pyramidal neurons, 50 μM Alexa-594 and 250 μM Fluo-4 (Invitrogen) were added to the pipette solution, as previously described ^17^. Alexa-594 fluorescence was used to reveal neuronal morphology, whereas fluorescence signals emitted by Fluo-4 were used for calcium imaging. L5 neurons with a long axon in the plane of the slice were selected. Laser sources for fluorescence excitation were set at 488 nm for Fluo-4 and 543 nm for Alexa-594. Emitted fluorescence was collected between 500 and 580 nm for Fluo-4 and between 620 and 750 nm for Alexa-594. After whole-cell access, the dyes were allowed to diffuse for at least 15-20 minutes before acquisition of fluorescence signals. Action potentials were evoked by a brief (2 ms) and strong (2-3 nA) pulse of depolarizing current with or without a preceding subthreshold depolarization (50 ms, 50-100 pA). Electrophysiological signals were synchronized with calcium imaging in line-scan mode (speed, 0.5-2 kHz). Acquired Fluo-4 signals were converted to ΔF/F values and peak amplitudes were measured with a custom-made analysis program (LabView, National Instruments).

### Axonal recordings

Simultaneous recordings from the soma and axonal cut ends (blebs, at 50-152 μm from the cell body) were obtained in L5 pyramidal neurons, as previously described ^17^. The axon was recorded 10-15 min after whole-cell access of the somatic compartment and the full labeling of the axon by Alexa 488 and visualized with the LSM-710 confocal microscope.

### Network activity

To test the incidence of spiking activity on cortical network activity, paired-recordings from adjacent L5 pyramidal neurons were obtained in physiological extracellular calcium conditions (1.3 mM). A spike was elicited in one of the two cells after having verified the integrity of the axon in the spiking neuron. For this, neurons were recorded with 50 μM Alexa 488 and only recordings in which the spiking cell displayed an axon longer than 100 μm were kept for final analysis. The spontaneous excitatory postsynaptic activity was measured in the non-spiking cell by counting the number of post-synaptic events occurring 200 ms before and 200 ms after the spike with the use of a custom-made software programmed in LabView. 6 out of 9 recordings were obtained from cells that were not connected by a synapse. The remaining 3 pairs were monosynaptically connected. As the results were qualitatively similar for the two groups, we concluded that the increase in spontaneous activity induced by a spike triggered by synchronous-like input was due to the enhancement of network activity and not to asynchronous release.

### Pharmacology

Tetrodoxin (TTX) and carbamazepine (CBZ) were provided by Tocris. All drugs were bath applied.

### Dynamic clamp

Excitatory postsynaptic conductance (**Fig. 1A**) was injected into the presynaptic neuron using a dynamic-clamp system (SM-1; Cambridge Conductance, Cambridge, UK) and a computer conductance profile delivered via an analog-digital interface (Digidata 1440A; Molecular Devices). The conductance profile was derived from the AlphaSynapse Point Process from the NEURON 7.4 software. The reversal potential for AMPA-receptor-like component was set at 0 mV.

### Hodgkin-Huxley modeling

A multi-compartment model of L5 pyramidal neuron was simulated with NEURON 7.4. The neuronal morphology was taken from Hu and coworkers ^20^. All simulations were run with 10-μs time steps and the nominal temperature of simulation was 33°C. The voltage dependence of activation and inactivation of Hodgkin-Huxley-based conductance models were taken from Hu et al. 2009 ^20^ for g_Nav1_._2_ and g_Nav1_._6_, from Mainen et al. 1995 ^41^ for g_KDR_, from Shu et al., 2007 ^42^ for g_Kv1_, and from Bischofberger et al, 2002 ^43^ for g_Ca_. The equilibrium potentials for Na^+^, K^+^, Ca^2+^ and passive channels were set to +60 mV, −90 mV, +140 mV and −70 mV, respectively. The conductance density is provided in **Supplementary Table 1**. The resting membrane potential was set to −78 mV. The current was injected in the soma to produce action potentials. EPSPs were simulated in **Fig. 5A** with synaptic conductance whose waveform was determined by 2 exponential functions (EPSC rise-time: 1.5 ms, and EPSC decay: 10 ms) and matched experimentally determined mEPSCs ^44^. In other modeling figures (**Fig. 5B**, **Supplementary Fig. 6**) action potentials evoked by synchronous-like inputs were produced by injection of 1.6 nA during 2 ms directly from the resting membrane potential. Action potentials evoked by asynchronous-like inputs were produced by injection of 1.6 nA during 2 ms after a 50 ms subthreshold depolarization via injection of 0.3 nA. In order to observe the postsynaptic response, a presynaptic terminal of the L5 neuron located at 150 μm from the soma and located on an axonal collateral was synaptically connected to a postsynaptic neuron. The postsynaptic neuron was a single compartment model containing only passive conductance. The level of release was dependent on the Ca^2+^ concentration in the presynaptic terminal. The synapse model was taken from Destexhe et al., 1994 ^45^.

### Network modelling

#### • Neuron model

Simulations were performed using current-based leaky integrate-and-fire neurons with membrane time constant *τ_m_* = 20 *ms* and resting membrane potential *V_rest_ =* −70 *mV*. When the membrane potential *V_m_* reaches the spiking *V_thresh_ =* −55 *mV*, a spike is generated and the membrane potential is clamped to the reset potential V_reset_ = −70 *mV* during a refractory period of duration *τ_ref_ =* 5 ms. Each presynaptic spike produces an exponentially decaying current with time constant *τ_exc_* = 5 *ms* for excitation and *τ_inh_ =* 10 *ms* for inhibition. All neurons are stimulated with a zero-mean white noise current ξ(*t*) resulting in voltage fluctuation of standard deviation *σ* = 4 *mV*. The model equations are thus:

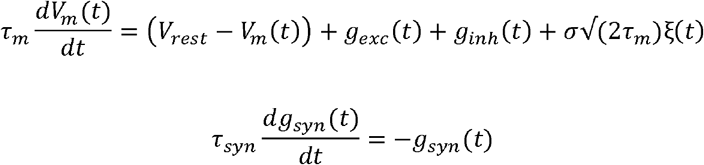

where *syn* ∈ {*exc, mh*} and each presynaptic spike triggers an instantaneous increase: *g_syn_ → g_syn_ + λ_syn_w_i_*, where *w_i_* is the synaptic weight and *λ_syn_* is a scaling factor calculated so that a synaptic weight of 1 mV produces PSPs of peak size 1 mV. Note that *g_syn_* is in units of volt, i.e., the membrane resistance is implicitly included in the variable. For exponential synapses, we can compute analytically:

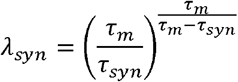

#### • Network model

We used a random balanced network ^46^ composed of 4000 excitatory neurons and 1000 inhibitory neurons. Every neuron in the network is connected to 10% of the others, with delays drawn from a uniform distribution between 0 and 5 ms. To set the network in a balanced regime, we neglected the contribution of spikes and used Campbell’s theorems, which give the mean and variance of a shot noise (corresponding to sums of PSPs with Poisson statistics):

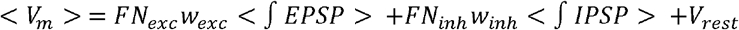

where *F* is in the firing rate of the network, and *N_syn_* the average number of incoming synapses per neuron. In the balanced regime, we want to have < *V_m_ > = V_rest_*, so we should have

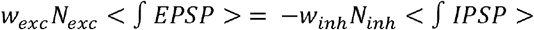

Since the connection probability is uniform, we have *N_exc_* = 4*N_inh_*. Using the fact that < *∫ EPSP* > = *λ_exc_τ_exc_* and < *∫ IPSP* > = *λ_inh_τ_inh_*, we have the following equation for the ^balance^ *g w_inh_/w_exc_*

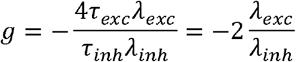

Initial synaptic weights for recurrent excitatory connections are drawn from a log-normal distribution such that < *w_exc_* > = 1 *mV*, in order to match experimental findings ^47^ (see **Supplementary Fig. 8** panel b and c). To be in a strong balanced regime, recurrent inhibitory connections are drawn from a Gaussian distribution such that < *w_inh_* > = −1.5g < *w_exc_* >. Standard deviation of the Gaussian distributions is chosen to be a fourth of the means.

In the spontaneous regime, the network displays an asynchronous irregular regime characterized by a low firing rate (0.2 Hz) (see **Supplementary Fig. 8** panel a, d and f), and an average coefficient of variation for the inter-spike intervals close to 1 (see **Supplementary Fig. 8** panel e).

#### • Protocol

To mimic the experimental protocol, we randomly selected 100 excitatory neurons within the network, and forced them to spikes at controlled times. In half of the trials (100 repeats), the spike had a “normal” effect (ai-AP), while in the other half of the trials (100 repeat), to mimic the fact that this particular spike was considered to be a si-AP spike, all the effective weights of the spiking neuron were artificially increased by 30%. Injections of consecutive spikes were spaced by a short duration of 200 ms, and the si-AP spike effect was limited to the effective weights onto excitatory neurons. Simulations where all effective weights were increased (both to excitatory and inhibitory neurons) showed that the effect was strongly reduced (data not shown).

#### • Simulator

All simulations were performed using the Brian simulator version 2 ^48^ with a fixed time step of 0.1 ms.

